# Human occipital bone sexual dimorphism in an Archiac period archaeological population

**DOI:** 10.1101/177758

**Authors:** Dr. Kara C. Hoover

## Abstract

**PREPRINT NOTES:** This paper was submitted for publication the *International Journal of Osteoarchaeology* in August of 2003. The paper was rejected roughly a year later due to scope and collection issues. I revisited the paper in 2007 with an eye to updating and resubmitting with additional data but the project was triaged from the publication pipeline due to the work involved relative to other projects more relevant to my research interests at the time. The current preprint paper contains updated references on occipital bone sexual dimorphism—all limited to the condylar region rather than the full suite of variation in the region.

I am posting this to the preprint server to contribute the findings to the larger scientific community in the hopes that I might have time to revisit the paper as a student project. A more rigorous study would need to expand the sample either to a study collection that has remains with known sex (either verified via molecular sexing techniques or using a modern study collection) or additional comparative collections from hunter-gatherer populations. If a student is interested in taking up the topic and assuming lead authorship on a co-authored paper, I’d be happy to collaborate and see this piece through the publication pipeline.

**ABSTRACT:** This paper presents the results of a comprehensive study of sexual dimorphism in the human occipital region of the skull using the remains of 39 prehistoric adults from the prehistoric Windover (6,000-8,000 BP). The results of a discriminant analysis classified 64% of the individuals correctly (using osteologically identified sex). Sexual dimorphism in this population may not be prevalent due to environmental stress.

## Introduction

The occipital lobe of the brain houses both the vision center and the area that controls voluntary skeletal movement (cerebellum). The evolutionary significance of the occipital lobe to both vision and bipedalism is corollary to the functional significance of the occipital bone. In fact, the heavy buttressing of the bone serves as both protection for the vision center and cerebellum as well as an attachment site for major neck muscles. As such, structural components of the bone may be sexually dimorphic in expression of various traits. Metric and nonmetric traits of the occipital are not typically reported although some attempts at standardization have been made (Golekon and Turgut, 2003; Olivier, 1969). Rather, features such as the external occipital protuberance (EOP) and muscular development are nominally described or rank-ordered. In addition, morphometric analyses of the condylar region have been conducted for clinical purposes (Naderi et al., 2005; Saluja et al., 2016).

Previous studies on sexual dimorphism in the human occipital region focused on radiographic analyses of selected features (Golekon and Turgut, 2003; Hsiao *et al.*, Lui, 1996), biomechanical analysis of spinal rotation relative to the condylar region (Chancey et al., 2007), and morphometric studies of the basal occipital region in cadavers (Kumar and Nagar, 2014; Varsha et al., 2017), study collections (Holland, 1986; Macaluso, 2011; Wescott and Moore-Jansen, 2001), and archaeological populations (Gapert et al., 2008; all find limited utility in metric sex identification using discriminant function analyses and recommend application of metric sex identification only in the case of a fragmented cranial base with no other methods available. Only one paper reports reliable metric identification but their data were collected using three-dimensional computed tomography—not a commonly available forensic or bioarchaeological tool (Uysal et al, 2005). The analysis of sexual dimorphism for the current study is more comprehensive in the use of endocranial and ectocranial metric and non-metric traits of the occipital region, not limited to the condylar area.

## Materials and Methods

The data for this study were collected from 39 well-preserved occipital bones of adults from the Windover site, a prehistoric Florida mortuary pond that dates from 8000 to 6000 BP. Skeletal materials are unusually well preserved due to the anaerobic environment of the bog in which they were deposited. Population demography is well balanced between the sexes, and individual ages range from infancy to 65+ (Purdy, 1991). The Windover population were food-foragers and likely engaged in gender-based divisions of labor which emphasized short bursts of intense activity for males and repetitive endurance activity in females. This differentiation of labor generally induces sexually dimorphic traits.

Ectocranial metric features consist of two measurements: depth of EOP and distance between the lambda and inion (LI). White (1991) locates the EOP on the ectocranial midline at the intersection between the occipital and nuchal planes. Olivier (1969) more specifically locates it on the inferior or superior nuchal line. Using Olivier's location, the EOP was measured for depth at the inion, or center of the EOP. LI distance is measured from the inion to the point where the lambdoidal suture meets the sagittal suture, the lambda (White, 1991; Olivier, 1969). Ectocranial nometrics also use standard landmarks of the skull (White, 1991) and include the presence or absence of a nuchal crest and number of associated nuchal lines. The general form of the EOP is based on Olivier’s classification of lateral skull shape (Olivier, 1969): flat, round, protuberant, or modified.

Endocranial features again use standard landmarks of the human skull (White, 1991) and were entirely non-metric. The location of the occipital or superior sagittal sulcus, which joins the transverse sinus and serves as the major vehicle of blood drainage from the brain to the body, was scored relative to the cruciform eminence as right, left, or indeterminate. Cases were indeterminate when the sulcus was only faintly expressed, not clearly discernible as the sulcus, or appeared as accessory to the cruciform eminence. The internal occipital protuberance (IOP), lying at the center of the endocranial cruciform eminence, was located relative to the EOP as either above, below, or concurrent.

All metric traits were measured with an electronic sliding caliber, with an average of three measurements to assess intra-observer error. Nonmetric data are rank-ordered and also an average of three observations. Discriminant function coefficients were calculated for the sample to test the utility of the proposed method of sex estimation.

## Results and Discussion

Basic descriptive statistics suggest that certain traits might be more dimorphic than others (Table 1). Females tend to exhibit greater variation around the mean than males in the expression of four of the seven traits: depth of EOP, LI, nuchal crest, and general form. This greater variation in females may cause an incorrect estimation where traits overlap with males. The IOP values for males and females are roughly equal. The general form of the occipital bone for both sexes was round. This is surprising as the male expression of this trait is expected to be protuberant due to more rigorous muscle attachments. Interestingly and contrary to the greater variation exhibited more frequently by females, the sinus sulcus has a consistent expression to the right side in females (67% versus 14% to the left) while expression in males is evenly distributed (Table 2).

**Table 1.** Descriptive Statistics.

|  |  | Sex |  |
| --- | --- | --- | --- |
|  |  | Male | Female |
| Depth of EOP | Mean | 5.6267 | 5.0000 |
|  | Std. Deviation | 1.5691 | 1.9992 |
| LI | Mean | 71.0067 | 67.8737 |
|  | Std. Deviation | 11.8542 | 12.1325 |
| Nuchal Crest | Mean | .6667 | .5263 |
|  | Std. Deviation | .4880 | .5130 |
| Nuchal Lines | Mean | 4.6000 | 4.2105 |
|  | Std. Deviation | 1.7238 | 1.3976 |
| General Form | Mean | 1.6000 | 1.8947 |
|  | Std. Deviation | .5071 | .5671 |
| IOP | Mean | 2.6000 | 2.5789 |
|  | Std. Deviation | .6325 | .6070 |
| Sulcus Site | Mean | 1.6667 | 2.0000 |
|  | Std. Deviation | .7237 | .5774 |

**Table 2.**
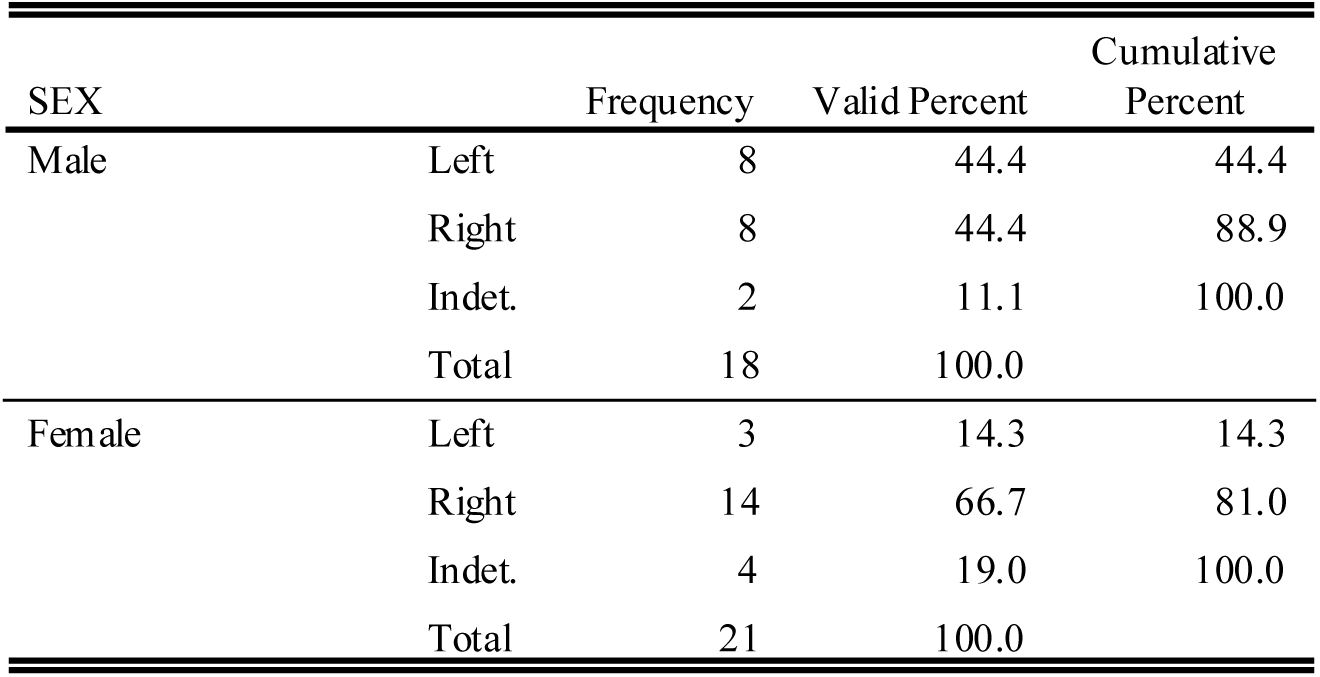
Sinus Sulcus Frequency.

Using SPSS, discriminant analysis of metric and nonmetric variables successfully grouped 90% of all individuals. Cross-validated results, however, grouped only 64% of the cases successfully (Table 3)—these results are similar to other studies using discriminate function on the occipital basal region (Holland 1986; Gapert et al, 2008; Macaluso, 2011; Uysal et al, 2005; Wescott and Moore-Jansen, 2001). The cross-validated classification results indicate the general efficacy of the function outside the data from which it was developed. Therefore, with regard to the cross-validated classification results, the proposed method was not useful for this sample. Coefficients with the heaviest weighting, and, therefore, most contributive to the model were depth of sinus sulcus, EOP, nuchal crest, and LI.

**Table 3.**
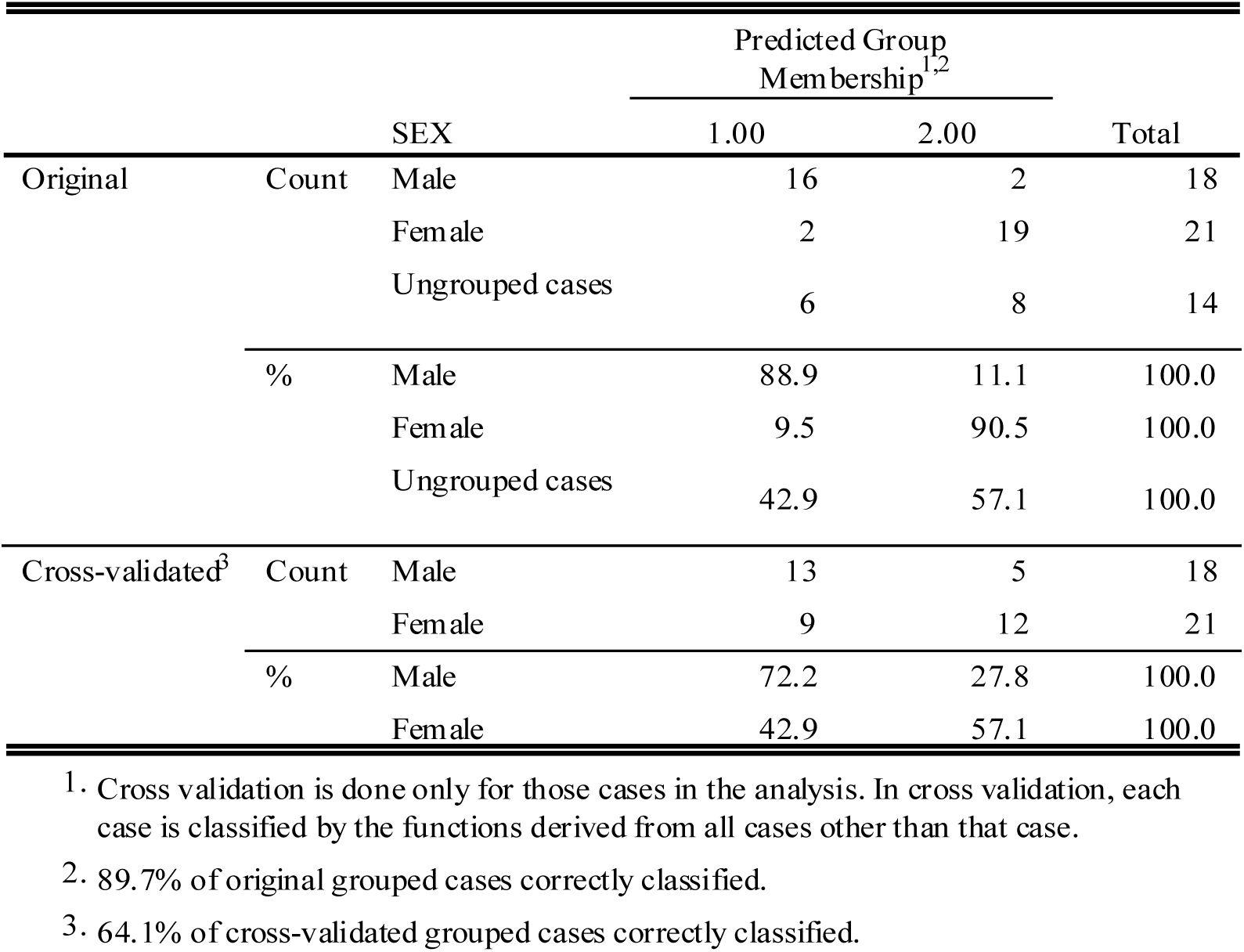
Classification Results of Discriminant Analysis.

**Table 4.**
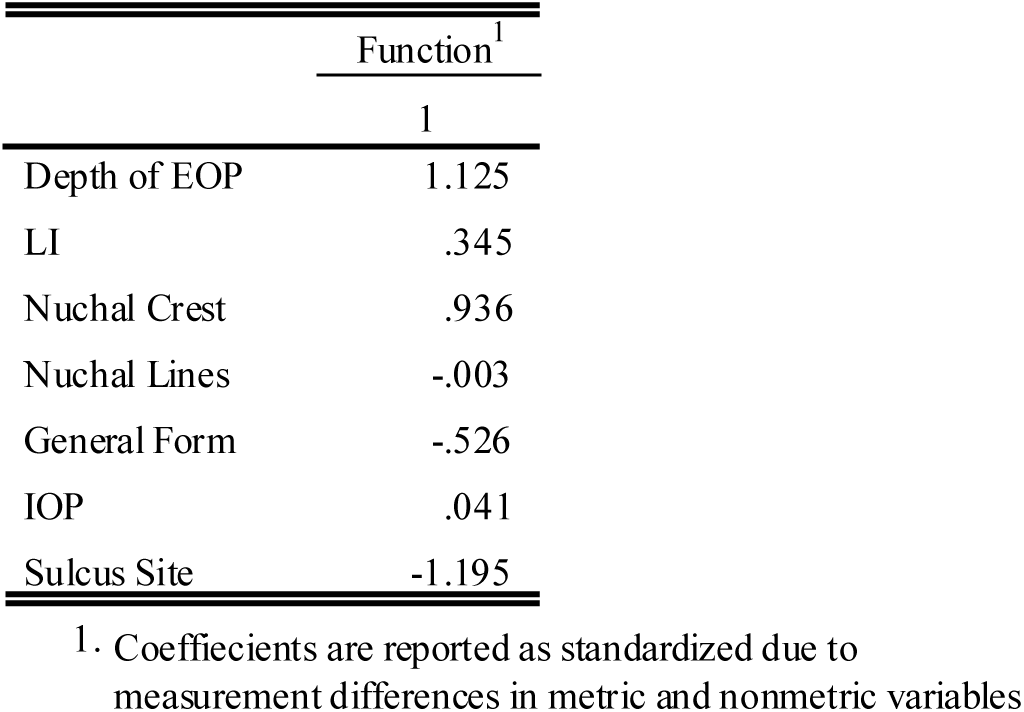
Canonical Discriminant Function Coefficients, Standardized.

|  | Function<sup>1</sup> |
| --- | --- |
|  | 1 |
| Depth of EOP | 1.125 |
| LI | .345 |
| Nuchal Crest | .936 |
| Nuchal Lines | -.003 |
| General Form | -.526 |
| IOP | .041 |
| Sulcus Site | -1.195 |

1. Coefficients are reported as standardized due to measurement differences in metric and nonmetric variables

A post-hoc test of variation was conducted to determine if an unidentified factor, such as age or environmental stress, affected results. Data was explored with a univariate analysis of variance. Results indicate that only 41% of the variance is explained by sexually dimorphic variables (Table 5). The other 59% must be explained by other factors. Based on the findings of previous studies on aging in the occipital bone (Wescott and Moore-Jansen, 2001; Tocheri and Molto, 2002; Cool, Hendrikz, and Wood, 1995), age seems an unlikely confounding factor.

**Table 5:** Variance Analysis.

| Dependent Variable: SEX |  |  |  |  |  |
| --- | --- | --- | --- | --- | --- |
| Source | Type III Sum of Squares | df | Mean Square | F | Sig. |
| Corrected Model | 3.47<sup>1</sup> | 7 | .50 | 2.62 | .03 |
| Intercept | 1.27 | 1 | 1.27 | 6.71 | .02 |
| Depth of EOP | 1.69 | 1 | 1.69 | 8.96 | .01 |
| LI | .24 | 1 | .24 | 1.25 | .27 |
| Nuchal Crest | 1.09 | 1 | 1.09 | 5.75 | .02 |
| Nuchal Lines | .00 | 1 | .00 | .00 | .99 |
| General Form | .52 | 1 | .52 | 2.77 | .11 |
| IOP | .00 | 1 | .00 | .02 | .90 |
| Sinus Sulcus | 1.92 | 1 | 1.92 | 10.14 | .00 |
| Error | 4.91 | 26 | .19 |  |  |
| Total | 91.00 | 34 |  |  |  |
| Corrected Total | 8.38 | 33 |  |  |  |

<sup>1</sup>. R Squared = .414 (Adjusted R Squared = .256)

The primary other factor that might explain variation in the variables is changing human-environment interactions that resulted from a new adaptation to the environment. During the late Archaic period in Florida prehistory, the environment was transforming due to climate change— mega fauna were becoming extinct, the coastline was stabilizing, and the southern pine forest was advancing (Powell, 1995; Milanich and Fairbanks, 1980). In fact, dietary analysis of the group associated with the Windover site suggests the environment forced a change in adaptation, evidenced by a marine-based diet with a heavy reliance on saltwater shellfish (Powell, 1995). The limitations of diet and environmental pressure suggest that resources were extremely limited; indeed, the population experience high prevalence of nutritional deficiency markers (dental enamel hypoplasias) suggesting higher levels of biological stress during development (Berbesque and Doran, 2008). Typically, chronic environmental stress not only hinders growth and development in subadults but also reduces adult sexual dimorphism.

## Conclusion

The confounding factor to high confidence in sex identification via metric methods might be the environmental effects of the Archaic period that created a larger range of overlap between males and unusually robust females. The promise of this method, and, by association, the dimorphism of the occipital bone, is supported by previous radiographic studies of cephalometric (Hsiao *et al*., 1996) 3DCT metric data (Uysal et al., 2005) that used discriminant analysis to successfully group individuals. The proposed method suggested for estimating sexual dimorphism in the occipital bone, therefore, shows promise but further studies are recommended to explore the data and attempt to account for the potentially confounding factor of environmental pressure.

## References

Berbesque JC, and Doran GH. 2008. Brief communication: physiological stress in the Florida Archaic-enamel hypoplasia and patterns of developmental insult in early North American hunter-gatherers. Am J Phys Anthropol 136(3):351–356.

Chancey VC, Ottaviano D, Myers BS, and Nightingale RW. 2007. A kinematic and anthropometric study of the upper cervical spine and the occipital condyles. Journal of biomechanics 40(9):1953–1959.

Cool SM, Hendrikz JK, and Wood WB. 1995. Microscopic Age Changes in the Human Occipital Bone. Journal of Forensic Sciences 40: 789–796.

Gapert R, Black S, and Last J. 2009. Sex determination from the occipital condyle: Discriminant function analysis in an Eighteenth and Nineteenth Century British sample. American Journal of Physical Anthropology 138(4):384–394.

Golekon IN and Turgut HB. 2003. The external occipital protuberance: can it be used as a criterion in the determination of sex? Journal of Forensic Sciences 48: 513–516.

Holland TD. 1986. Sex determination of fragmentary crania by analysis of the cranial base. American Journal of Physical Anthropology 70(2):203–208.

Hsiao T-H, Chang H-P, and Liu K-M. 1996. Sex Determination by Discriminant Function Analysis of Lateral Radiographic Cephalometry. Journal of Forensic Sciences 41: 792–795.

Kunar A, and Nagar M. 2014. Human adult occipital condyles: a morphometric analysis. Research & Reviews: Journal of Medical and Health Sciences 3(4):112–116.

Macaluso PJ. 2011. Détermination biométrique du sexe dans une population française à partir de mesures de la région basale de l'os occipital. Bulletins et mémoires de la Société d'anthropologie de Paris 23(1):19–26.

Milanich JT and Fairbanks CH. 1980. Florida Archaeology. Academic Press: London.

Naderi S, Korman E, Citak G, Guvencer M, Arman C, Senoglu M, Tetik S, and Arda MN. 2005. Morphometric analysis of human occipital condyle. Clinical neurology and neurosurgery 107(3):191–199.

Olivier G. 1969. Practical Anthropology. C.C. Thomas: Springfield.

Powell JF. 1995. Dental Variation and Biological Affinity among Middle Holocene Human Populations in North America. PhD Dissertation. Texas A&M University: College Station.

Purdy BA. 1991. Windover. In The Art and Archaeology of Florida's Wetlands., Purdy BA (ed). CRC Press: Boca Raton; 205–228.

Relethford JH and Hodges DC. 1985. A Statistical Test for Differences in Sexual Dimorphism Between Populations. American Journal of Physical Anthropology. 66: 55–61.

Saluja S, Das SS, and Vasudeva N. 2016. Morphometric Analysis of the Occipital Condyle and Its Surgical Importance. Journal of clinical and diagnostic research : JCDR 10(11):AC01– AC04.

Tocheri MW and Molto JE. 2002. Aging fetal and juvenile skeletons from Roman Period Egypt using basiocciput osteometrics. International Journal of Osteoarchaeology. 12: 356–363.

Uysal S, Gokharman D, Kacar M, Tuncbilek I, and Kosa U. 2005. Estimation of sex by 3D CT measurements of the foramen magnum. Journal of forensic sciences 50(6):1310–1314.

Wescott DJ, and Moore-Jansen PH. 2001. Metric variation in the human occipital bone: forensic anthropological applications. Journal of forensic sciences 46(5):1159–1163.

White TD. 1991. Human Osteology. Harcourt, Brace, and Jovanovich: London.

